# Direct Measurement of Synchronous Precursor Selection (SPS) Accuracy in Public Proteomics Datasets

**DOI:** 10.1101/647917

**Authors:** Conor Jenkins, Aimee Rinas, Ben Orsburn

## Abstract

Reporter ion quantification techniques utilizing reagents such as TMT and iTRAQ allow proteomics studies to multiplex up to 11 different samples within a single LC-MS/MS experimental run. In these experiments, peptides derived from different samples are labeled with chemical tags possessing identical mass but differing distributions of heavy isotopes through their structure. Peptides from all samples may then be physically combined prior to LC-MS/MS. Relative quantification of the peptides from each sample is obtained from the liberation of low mass reporter ions alone, as these are the only discernible factor between peptides in the entire LC-MS/MS workflow. When coeluting ions of similar mass to charge ratios are fragmented along with the ions of interest, it is not possible to determine the source of the reporter fragments and quantification is skewed, most often resulting in ratio suppression. One technique for combatting ratio suppression is the selection of MS2 fragment ions that are likely to retain the intact mass tag region by synchronous precursor selection (SPS) and the liberation of the reporter ions from this combination of ions in MS/MS/MS (MS3). In this study we utilize a new post processing tool that can directly assess the accuracy of the SPS system for picking ions for quantification that are truly derived from the peptide of interest. We then apply this tool to the re-analysis of 3 public proteomics datasets. Directly assessing SPS accuracy allows a new measurement of confidence in the quantification values obtained from these reporter ion quantification experiments.

**Abstract Graphic:** 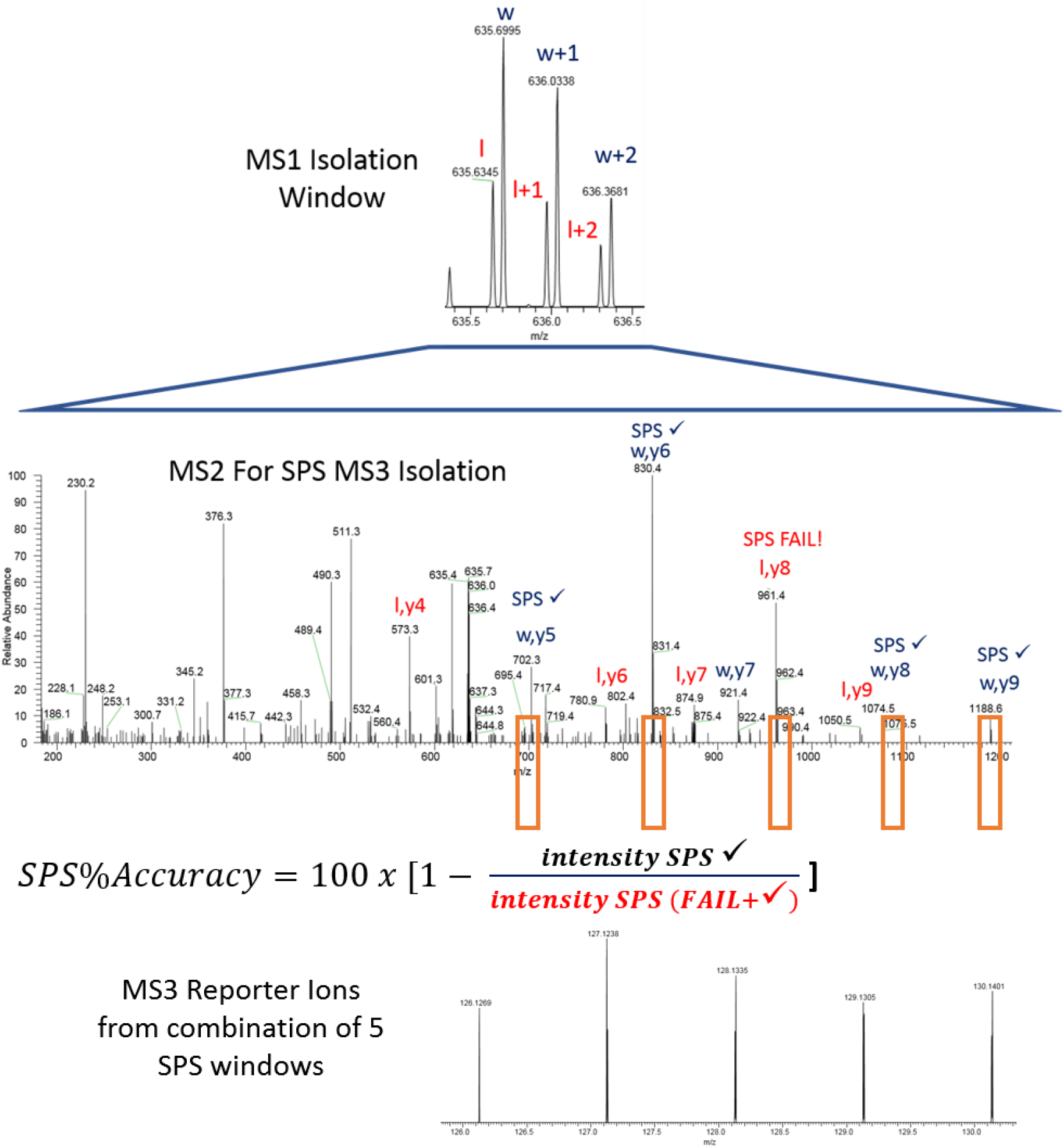

## Introduction

Reporter based quantification methods such as TMT and iTRAQ are nearly ubiquitous in proteomics labs world-wide. The ability to multiplex proteomics samples is an attractive alternative to label free quantification for several reasons. The first is the ability to increase instrument throughput by 11-fold with little decrease in depth of coverage. In addition, common instrument fluctuations such as spray instability are less impactful when all the peptides during the incident are affected identically. Reporter quantification does have a few well-documented downsides, most notably suppression of differential quantification values. In a complex mixture, many ions elute with ions of extremely similar mass to charge (m/z) ratios. If a peptide of interest has, for example, a true ratio of 5:1, but is fragmented with multiple coeluting ions with a ratio approximating 1:1, the combination of the released reporter ions from the coeluting species may result in an identification of the first peptide, but a ratio much lower than true for that identified peptide ^1^.

One popular mechanism for combatting reporter ion suppression is intelligent selection of MS/MS ions that remain in possession of their isobaric tags for MS3 based quantification. This was first developed by the selection of single high mass fragments for MS3 on a hybrid linear ion trap Orbitrap (LTQ-Orbitrap) system^2^. The ion selected from the MS2 was isolated in the ion trap and subjected to higher energy dissociation (HCD) to fully liberate the reporter ions. This method showed remarkable decreases in the obvious quantification suppression. Improvements in system architecture available on “Tribrid” systems such as the Orbitrap Fusion I and II devices have allowed “MultiNotch”^3^ or “Synchronous Precursor Selection” (SPS) where as many as 10 fragment ions may be selected from MS2 scans for HCD fragmentation resulting in additive liberation of the reporter ions ^4^. This selection is carried out in real time with ions selected that obey rules defined within the instrument parameters. Many indirect comparisons of SPS MS3 to MS2 based quantification of reporter ion experiments have been reported, including the recent description of a model system with completely missing proteins as a standard^5^. However, we do not believe a direct measurement of the accuracy of the SPS system has been reported. A new feature present in Proteome Discoverer™ 2.3 from Thermo Fisher Scientific allows the post processing comparison of peptide spectral match fragment ions to the extracted ion list used for SPS. In this study, we re-evaluate studies that employed SPS MS3 and assess the relative accuracy of SPS for each ion in selecting peptides for reporter liberation that are from the peptide of interest.

## Methods

### Files and Reanalysis

All RAW files were obtained from PRIDE via the ProteomeXchange web interface. Table 1 is a summary of the files obtained. FASTA sequences were derived from the UniProt/SwissProt annotated complete FASTA obtained 7/11/18 and parsed on the genus/species of the organism with IMP Proteome Discoverer 2.1 (www.pd-nodes.org). All analysis was performed in Proteome Discoverer 2.3 which was downloaded from the Thermo Flexera website on 3/7/19. All nodes and settings are described in Table 2. For assessment of SPS accuracy, 10 bins were made by adjusting the SPS % intensity parameter in the Consensus step. No other modifications were made to the sample processing workflows. Downstream data visualization was performed utilizing Proteome Discoverer 2.3 and the programming language R (version 3.5.2) along with the ggplot and ggpubr packages. The dplyr package was also used for data forming. All the code is made available online at https://github.com/jenkinsc11/sps_vs_Isolation.

**Table 1.** The files analyzed in this study

| Proteome Exchange ID | Instrument | Description of files analyzed | Reference |
| --- | --- | --- | --- |
| PXD005361 | Orbitrap Fusion | 2D fractionated human cell lines | Paulo and Gygi<sup>6</sup> |
| PXD004093 | Orbitrap Fusion | Single shot yeast digest | Liu *et al.*,<sup>7</sup> |
| PXD007145 | Fusion II "Lumos" | Single shot phosphoproteomics | Hogrebe *et al.*,<sup>8</sup> |

**Table 2.**
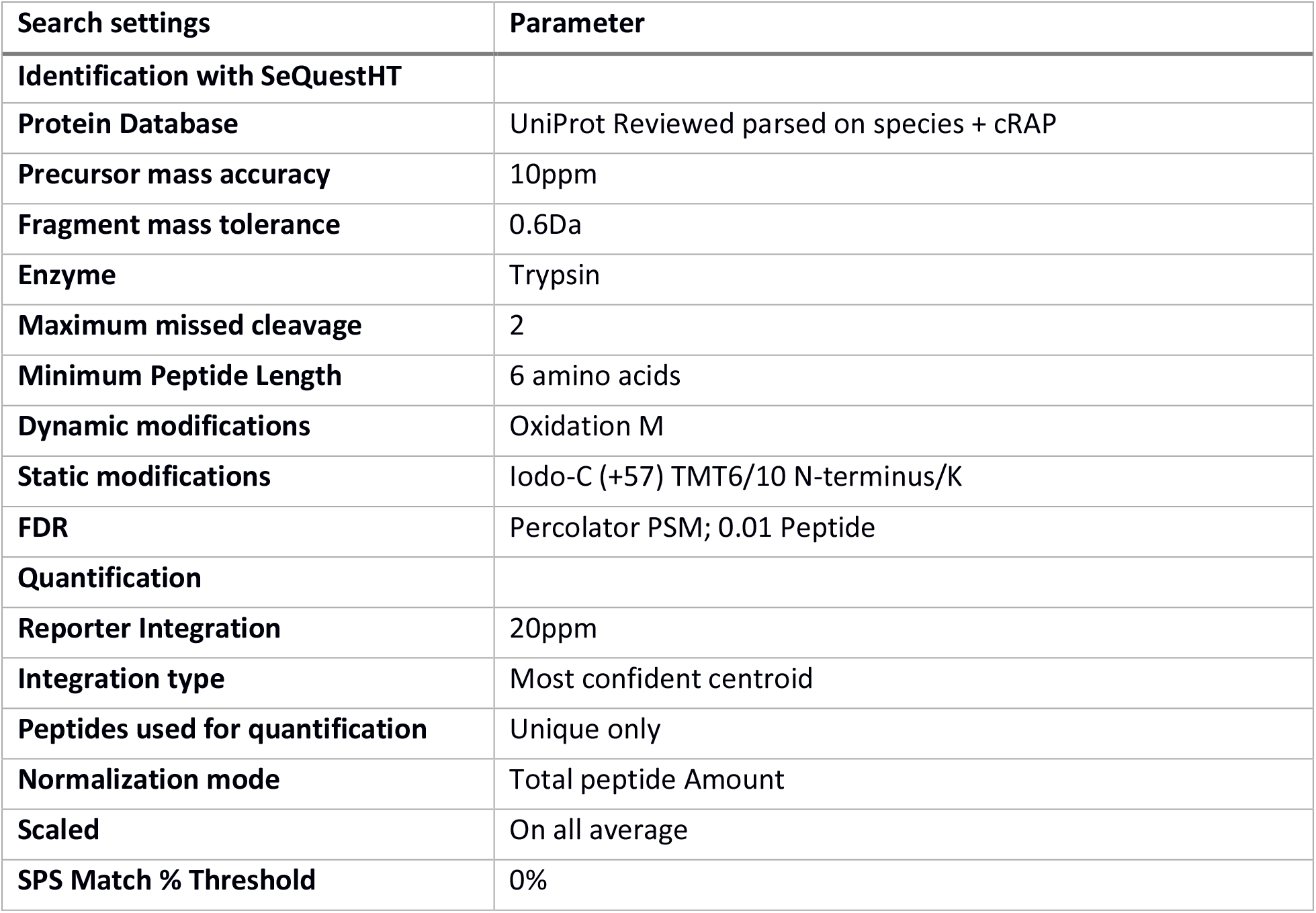
Search and quantification settings used for the study unless otherwise noted in results and results and discussions section.

| Search settings | Parameter |
| --- | --- |
| **Identification with SeQuestHT** |  |
| Protein Database | UniProt Reviewed parsed on species + cRAP |
| Precursor mass accuracy | 10ppm |
| Fragment mass tolerance | 0.6Da |
| Enzyme | Trypsin |
| Maximum missed cleavage | 2 |
| Minimum Peptide Length | 6 amino acids |
| Dynamic modifications | Oxidation M |
| Static modifications | Iodo-C (+57) TMT6/10 N-terminus/K |
| FDR | Percolator PSM; 0.01 Peptide |
| **Quantification** |  |
| Reporter Integration | 20ppm |
| Integration type | Most confident centroid |
| Peptides used for quantification | Unique only |
| Normalization mode | Total peptide Amount |
| Scaled | On all average |
| SPS Match % Threshold | 0% |

## Results and Discussion

Proteome Discoverer 2.3 contains a new feature for the measurement for matching the ions selected by SPS for MS3 quantification against the fragment ions used for each individual peptide spectral match (PSM). Re-evaluation of the SPS selection accuracy by automated consensus reports resulted in the number of retained peptides described in Table 2. Matching the fragments used for SPS to each PSM is relatively straight-forward. The simplest example is depicted by a cartoon in Figure 1 where a peptide possesses a single tandem mass tag. All b ions will lack the entire mass tag region and selection of these fragment ions by SPS will result in a decrease in the “SPS%Accuracy”. Ions resulting from multiple fragmentation events, such as the loss of the reporter ion region of the tag are easily excluded as many search engines cannot natively interpret ions resulting from more than one fragmentation event.

**Figure 1.**
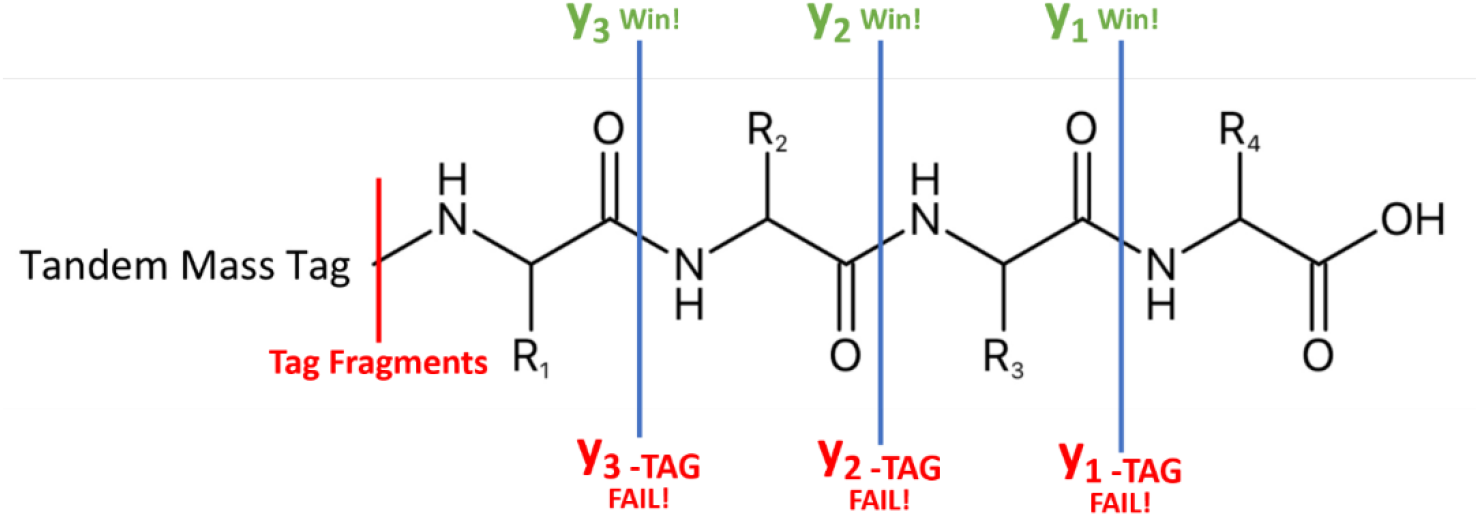
A cartoon demonstrating y ions that would be considered as correct SPS assignment compared to incorrect SPS assignment

**Figure 2.**
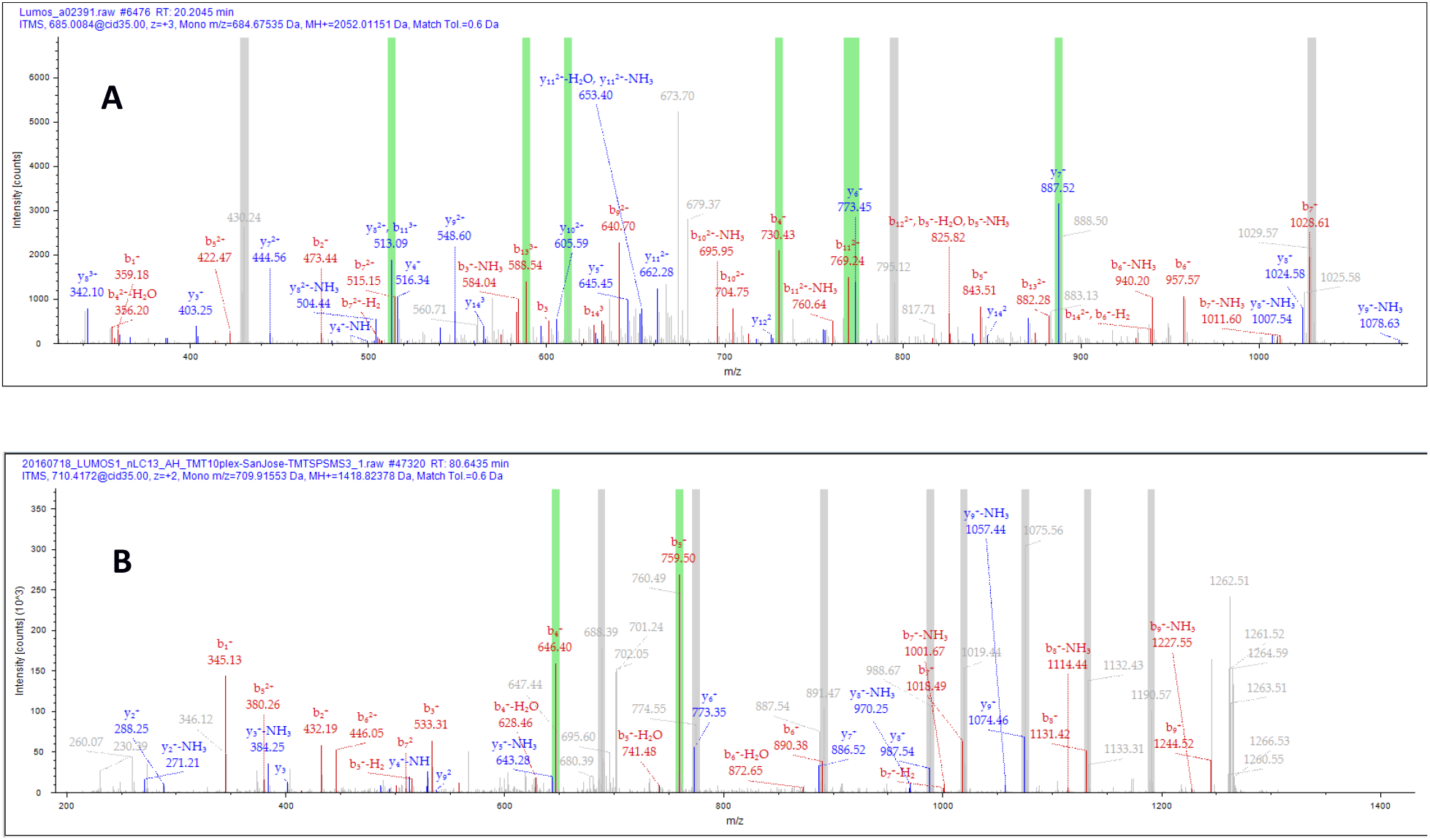
Representative PSMs highlighting the ions selected for SPS MS3 fragmentation. The green bars represent ions that were selected that could be attributed to the peptide of interest. Grey bars represent ions selected that could not be attributed to the peptide identified. A) A PSM with approximately 70% SPS precision. B) A PSM with a low %SPSAccuracy.

Verification of the accuracy of these outputs could be confirmed by manual inspection of a large percentage of labeled MS/MS spectra in Proteome Discoverer 2.3 (data not shown)

### Fractionated human cell lysates on Fusion generation I instrument (PXD005361)

A recent study form Paulo and Gygi^6^ studied the effects of the nicotine across 4 human cell lines. The peptides were TMT labeled, separated into 96 high pH reversed phase fractions and concatenated. The 12 resulting fractions were separated by a 180 minute gradient on a 30cm C-18 column. The authors report quantification of 8590 proteins across all cell lines. Our data processing pipeline produces highly similar results, with 8556 protein groups with quantification in all cell lines. (data not shown).

To visualize the accuracy of SPS in this set of 132,795 PSMs we have displayed the percentage of SPS signal matching fragments into 10 bins as shown in Figure 4. Over half of all PSMs (52.82%) were quantified from SPS selected ions that made up over 90% of all reporter signal. However, only 70.98% of all PSMs were quantified based on SPS fragments making up over 70% of the total reporter ion signal. The remaining 29.02% of PSMs were quantified based on SPS fragment ions that do not appear to be fragments from the peptide of interest.

**Figure 3.**
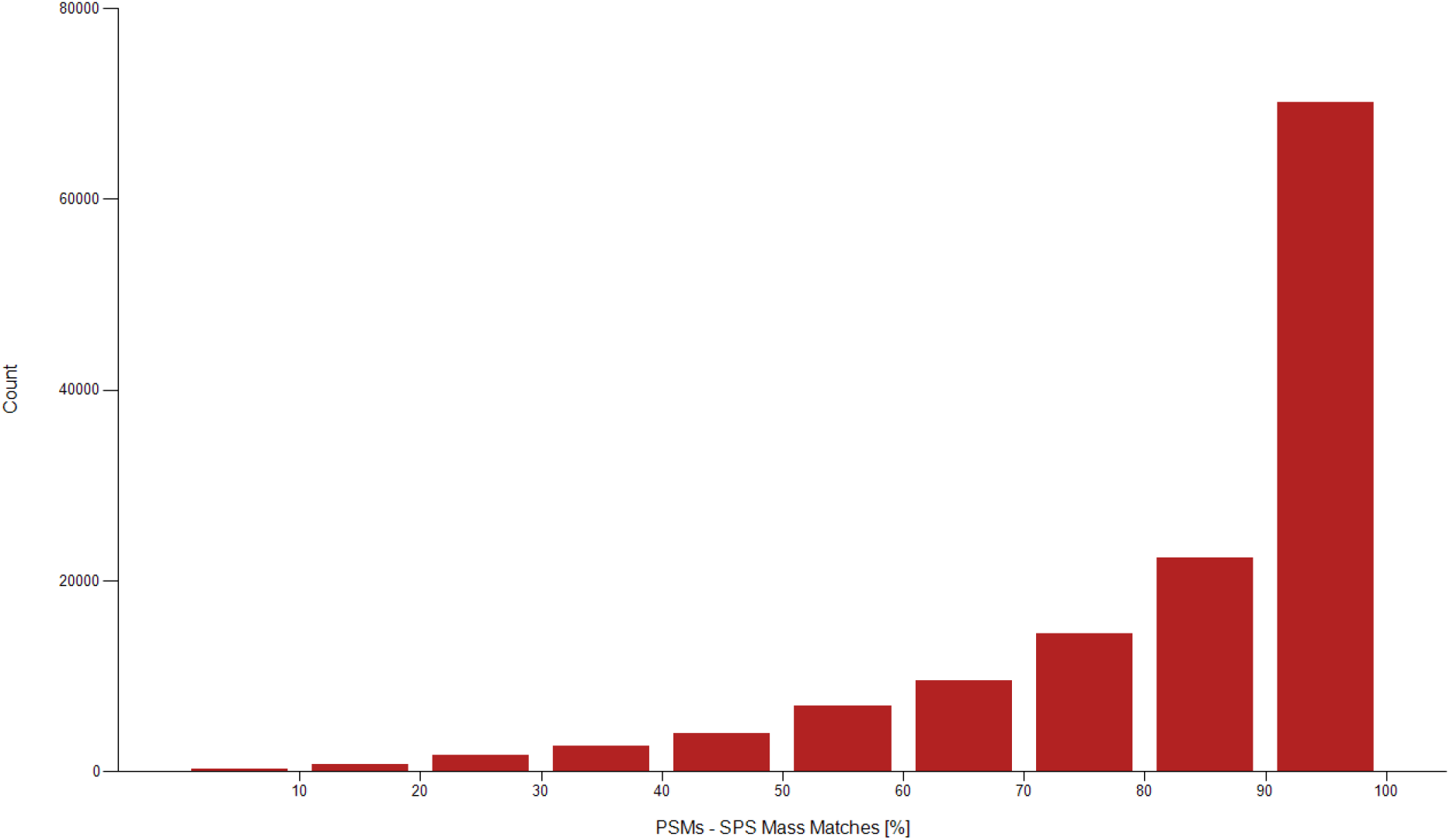
A histogram demonstrating the SPS match accuracy in 10% bins for TMT analysis of cell lines.

### Single shot medium complexity matrix (PXD004093)

A recent technical note from Liu *et al*., evaluated alternative methods for obtaining reporter ion quantification using the SPS methodology. With unit resolved TMT 6-plex reagent, this team compared the efficacy of reporter ion quantification where the MS3 was acquired in the linear ion trap following SPS. The samples were evaluated with two methods, the first being MS3 in the Orbitrap and identical experiments where the MS3 was acquired in the ion trap. On a digested and labeled mixture of *Saccharomyces cerevisiae* digests, acquiring quantification in the ion trap was shown to acquire more quantifiable peptides and proteins due to the higher relative speed and sensitivity of the ion trap and with no obvious changes in accuracy. The unfractionated samples were separated on a 20cm × 50µm ReproSil-Pur AQ C-18 column with a total run time of 130 minutes.

The results reported by the authors were consistent in our analysis with ∼2700 proteins quantified in each MS3 IT file and ∼2,100 proteins in each MS3 OT file. Previous studies have defined the detectable proteome of digests of this organism at approximately 4,000 proteins, all of which may be detected in a single shot analysis^9^. In order to integrate the reporter ions measured within the ion trap, the tolerance of the peak integration was widened to 0.4 Da. Wider tolerances were utilized, but no improvements in quantification appeared above this integration width (data not shown). No other alterations to the data processing pipeline was required.

As demonstrated in Figure 3, the SPS%Accuracy is highly similar whether the ion trap or Orbitrap is used for quantification of the reporter ions. The SPS%Accuracy appears similar to that of the highly fractionated human lysate. Approximately 50% of all ions quantified were based on measurements where SPS % accuracy was over 90%. In both sets of experiments approximately 75% of quantified PSMs were obtained from MS3 signal where SPS % accuracy was 70% or more.

**Figure 4.**
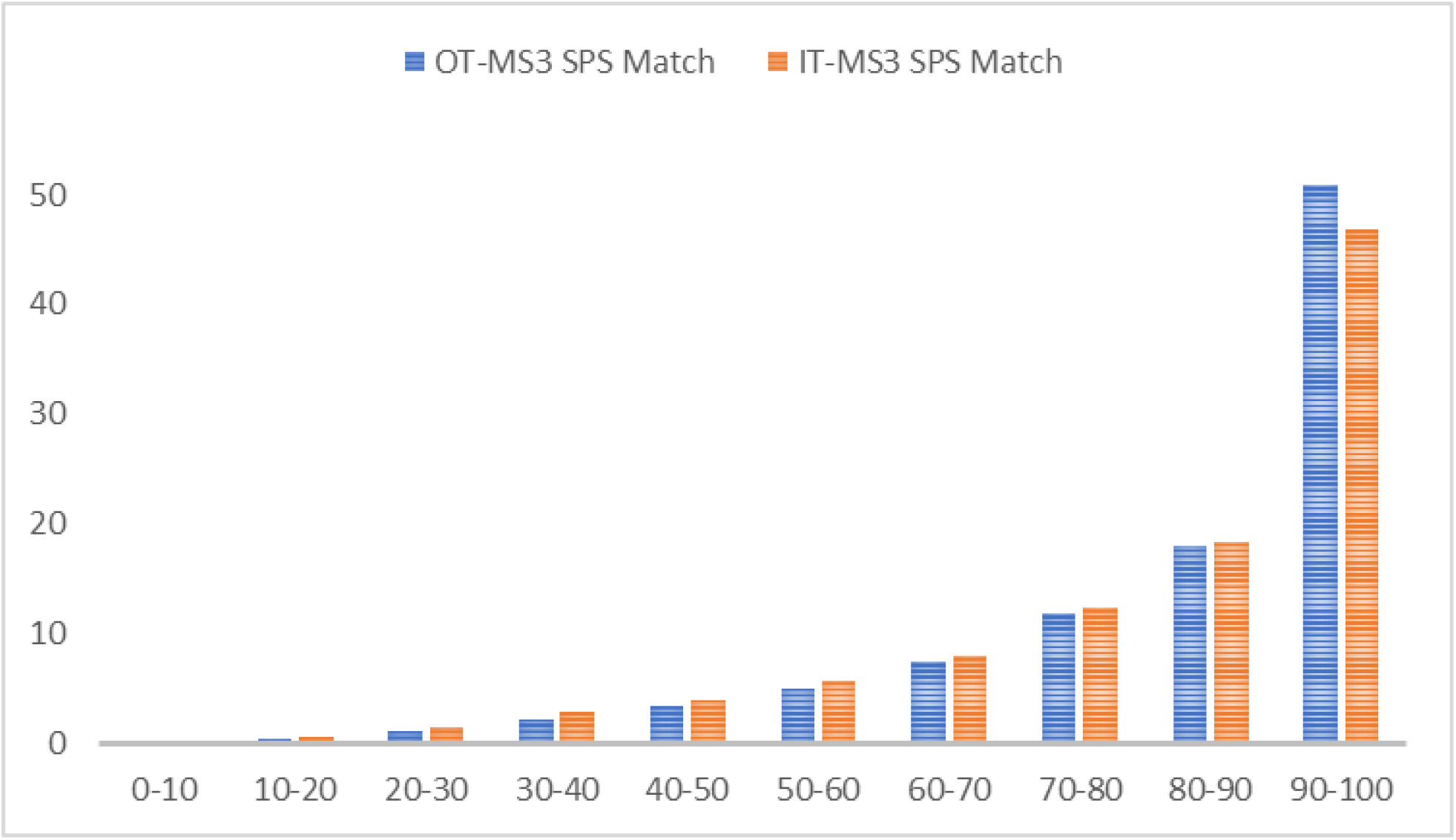
The SPS match percentage of the TMT 6-plex labeled yeast digests demonstrates little change whether the SPS selected ions are collected for quantification in the ion trap or Orbitrap detectors.

### Unfractionated Phosphoproteomics (PXD007145)

A recent study from the Olsen lab sought to compare common instrument methods for quantitative phosphoproteomics.^8^ The experimental design relied on a combination of enriched human and yeast digests. Three files publicly deposited utilized TMT MS3 according to manufacturer recommendations as the “San Jose” method files. These files were searched as described in Table 2, with the only alteration being the addition of the dynamic modification +79.966 on S, T, and Y.

The lack of prefractionation, even after phosphopeptide enrichment resulted in a complex sample and a corresponding reduction in the accuracy of the SPS selection process. Only 10.69% of PSMs had >90% SPS signal resulting from matched fragments. 36.63% achieved less than 50% SPS accuracy. A bar chart summarizing these results is shown in Figure 5.

**Figure 5.**
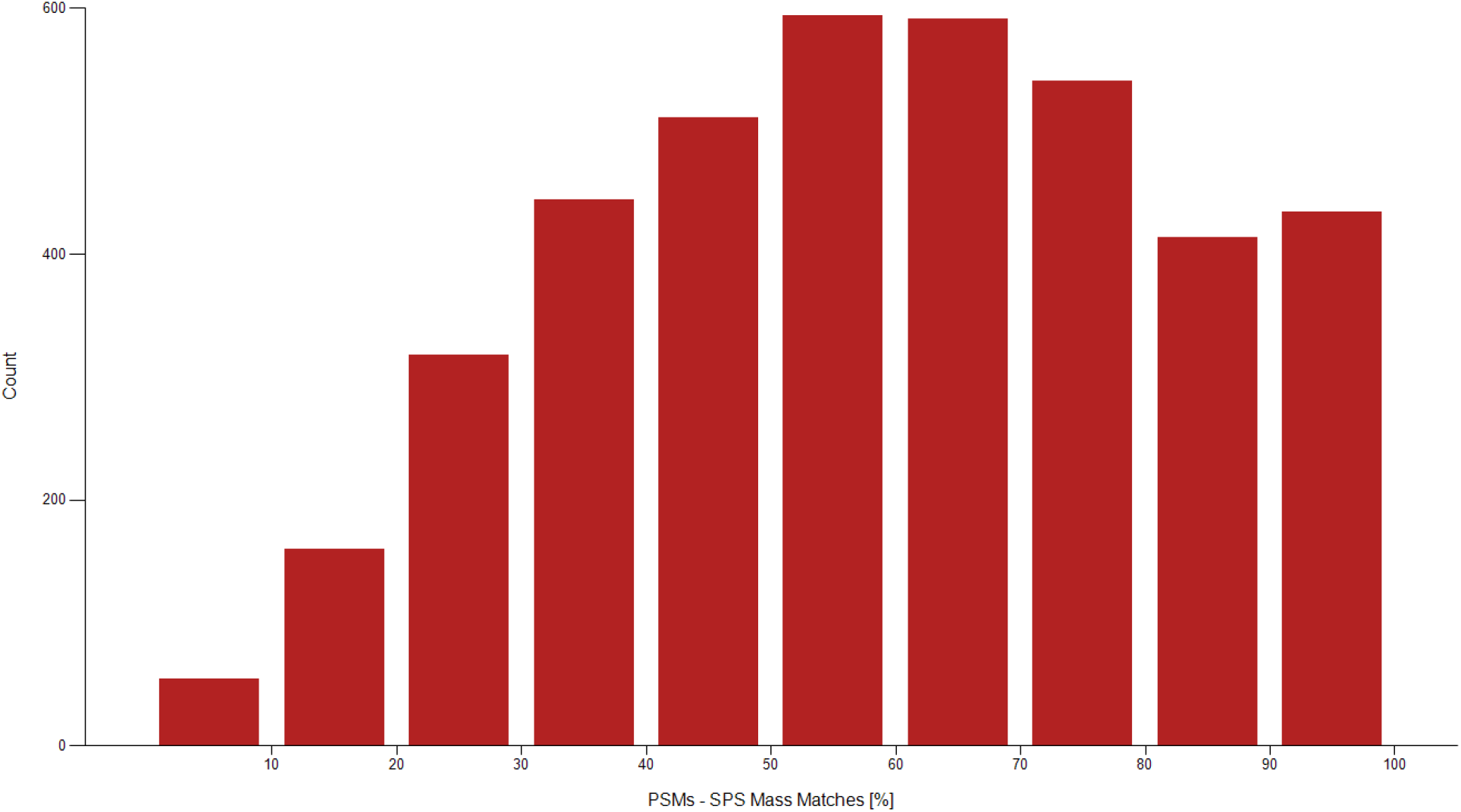
A histogram demonstrating the distribution of SPS MS3 matches from a reprocessed and unfractionated phosphoproteomics study.

### Comparing SPS Accuracy to Parent Ion Coisolation Interference

In order to determine the relative effects of parent ion coisolation interference on SPS%Accuracy, both values were extracted from the Proteome Discoverer output files for the PSMs from PXD005361. An example of the visualization of these data is shown in Figure 6. When these data are binned into 20%intervals it appears that most ions have both low isolation interference and high SPS%Accuracy. From this observation, we hypothesized that these two measurements would be correlative, but this visualization does not suggest this is the case. No direct correlation was found between the %Isolation interference and %SPS Accuracy in these three data sets analyzed (Supplemental Figure 2), suggesting that the two values are completely independent in the PXD005361.

**Figure 6.**
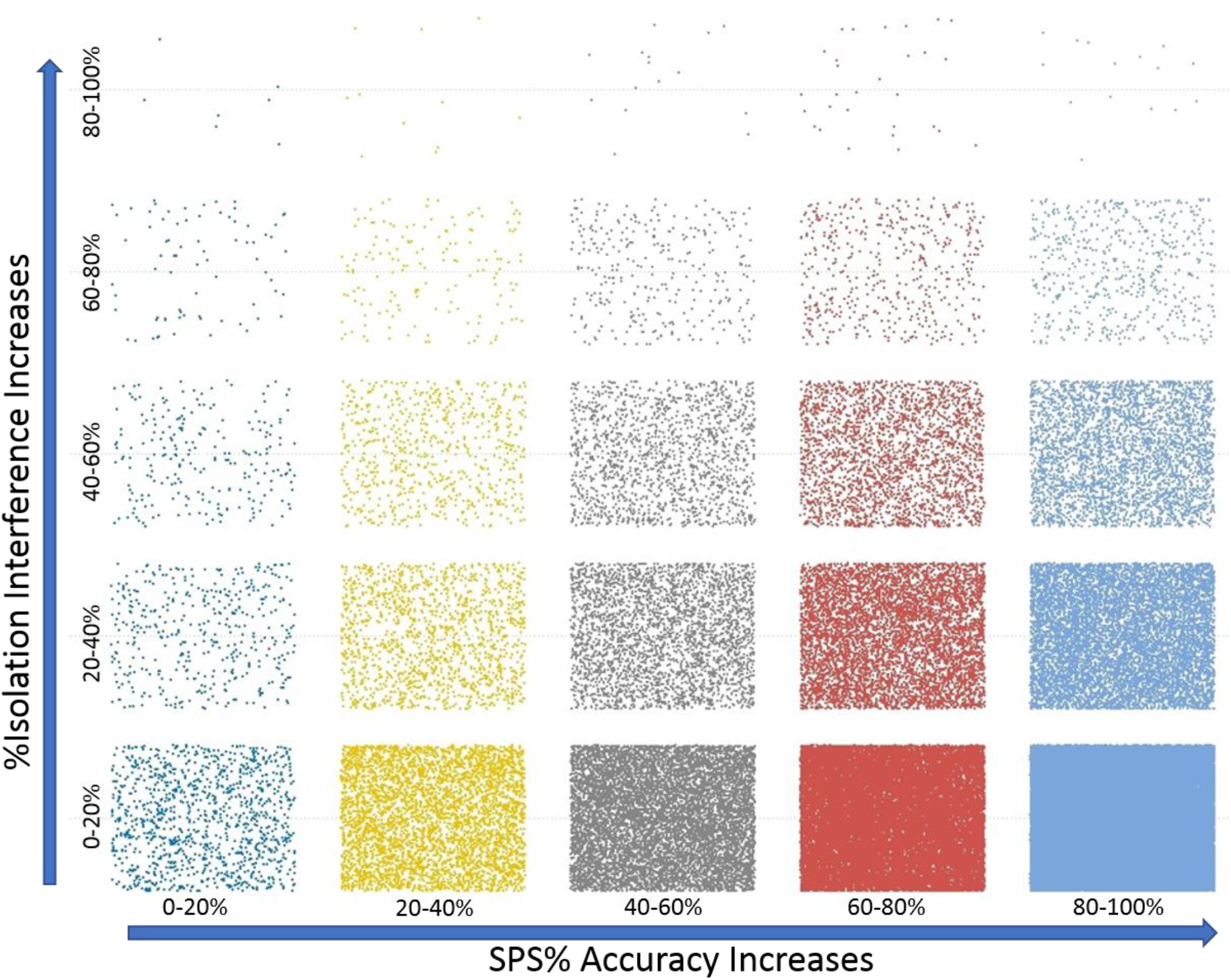
Binning of the % isolation interference and SPS% Accuracy by 20% relative cutoffs.

## Conclusions

We demonstrate a method whereby the accuracy of synchronous precursor selection can be estimated for any proteomics experiment. It is now common practice in MS2-based reporter ion quantification to discard ions with extremely high levels of coisolation, as the quantification will more closely match that of the coisolated ions than the ion of interest. Likewise, this measurement of SPS accuracy can be used as a filter, allowing ions with high degrees of mistaken identities within the SPS method to be eliminated from quantification. By reanalyzing 3 publicly available datasets we find that SPS is most accurate in less complex matrices or in highly fractionated samples. When complex samples are analyzed by single dimensional separation, the number of coeluting ions leads to correspondingly low SPS accuracy. Surprisingly, however, the SPS%Accuracy does not appear to directly correlate with the level of measured coisolation at of the MS2 selected ion.

It is worth noting that while we performed these analyses in Proteome Discoverer, the identities of the SPS selected windows for each MS3 scan can be extracted in alternative ways. The SPS windows for each MS3 event are listed within the scan header of the RAW files as shown in Supplemental Figure 1. This data can be extracted from the RAW file using Xcalibur and should be easily adaptable to other common data processing pipelines.

## Acknowledgements

We would like to thank Dr. Alexander Hogrebe for access to the TMT phosphoproteomic files. We would also like to thank Dr. Bernard Delanghe for helpful discussions and clarifications critical to the design of these studies.

---

**Supplemental Figure 1.**
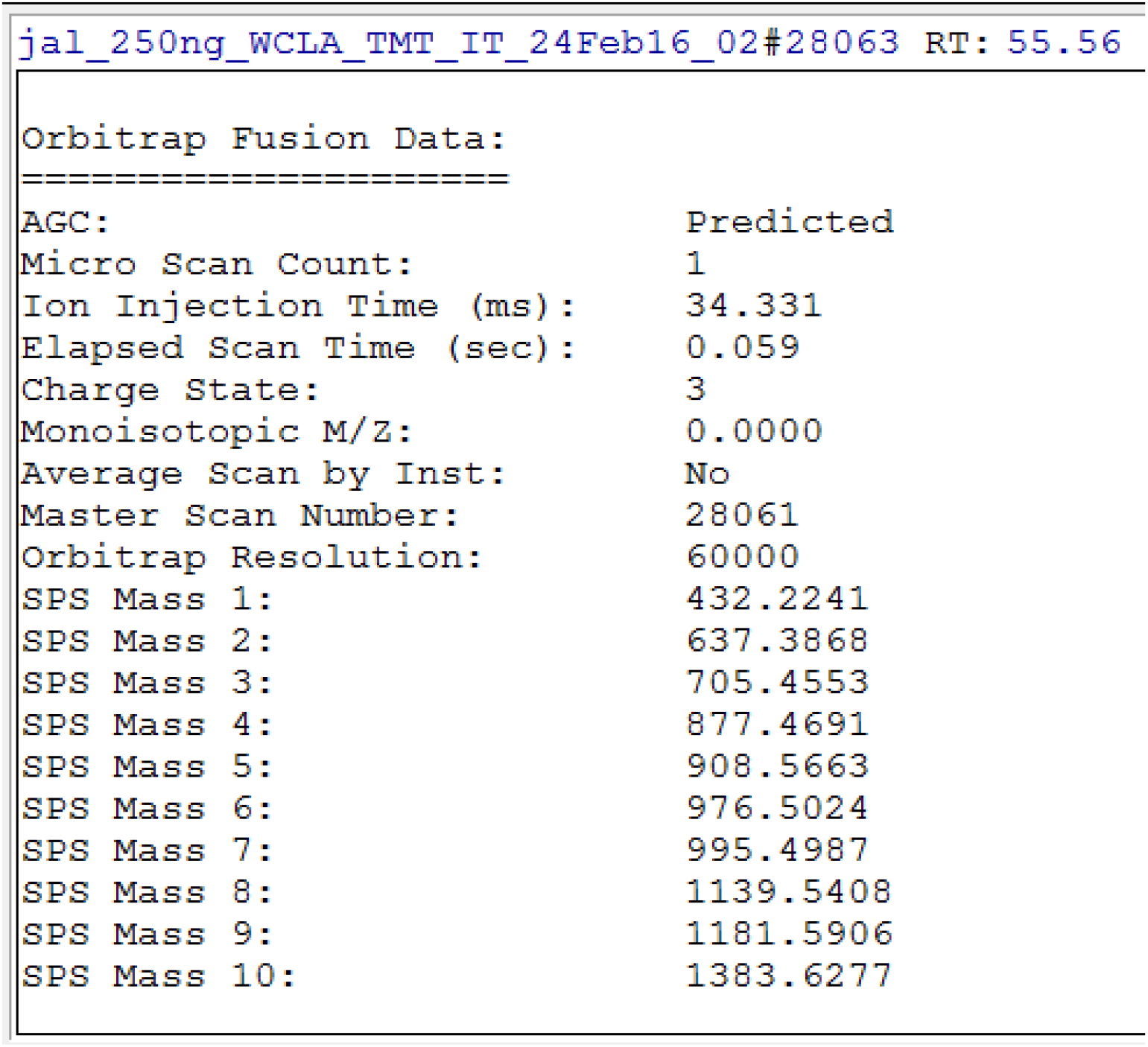
The location of SPS ions used for MS3 acquisition as visualized within the XCalibur software Scan Header.

**Supplemental Figure 2.**
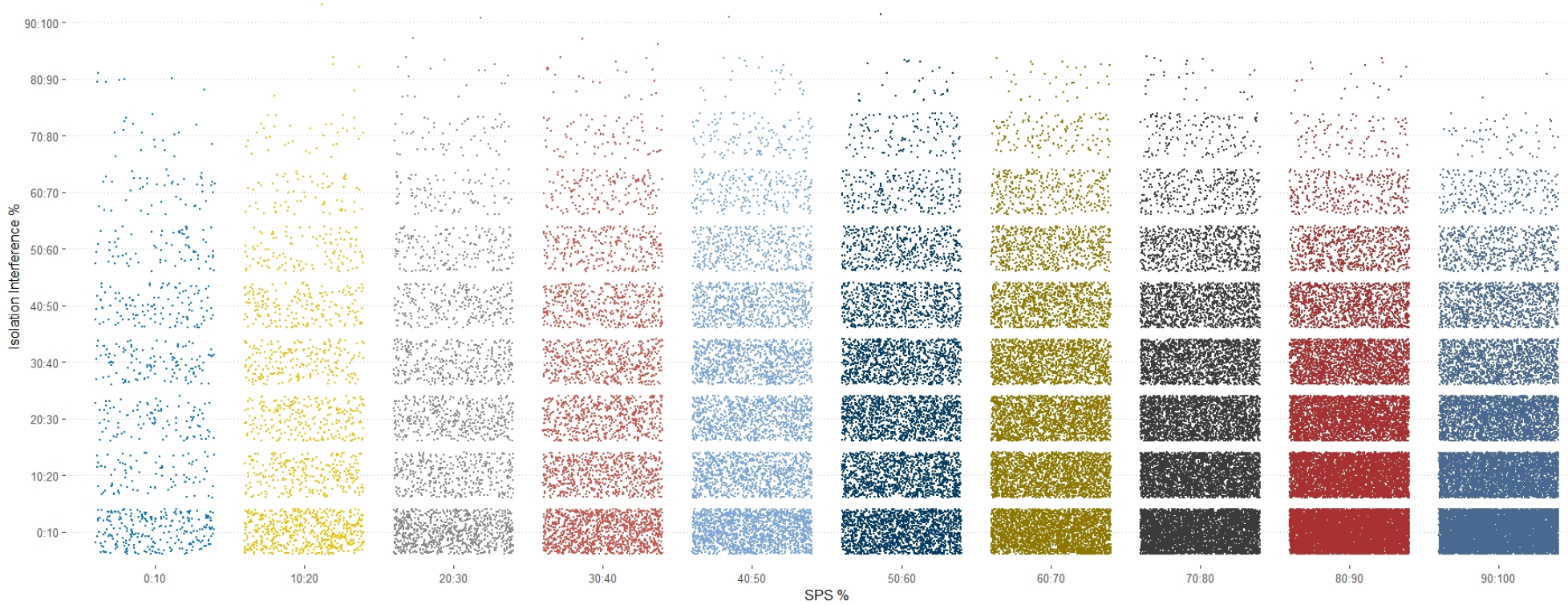
Binning of coisolation interference versus SPS%Accuracy of the Fusion Lumos files from PXD005361 in bins of 10.

**Supplemental Figure 3.**
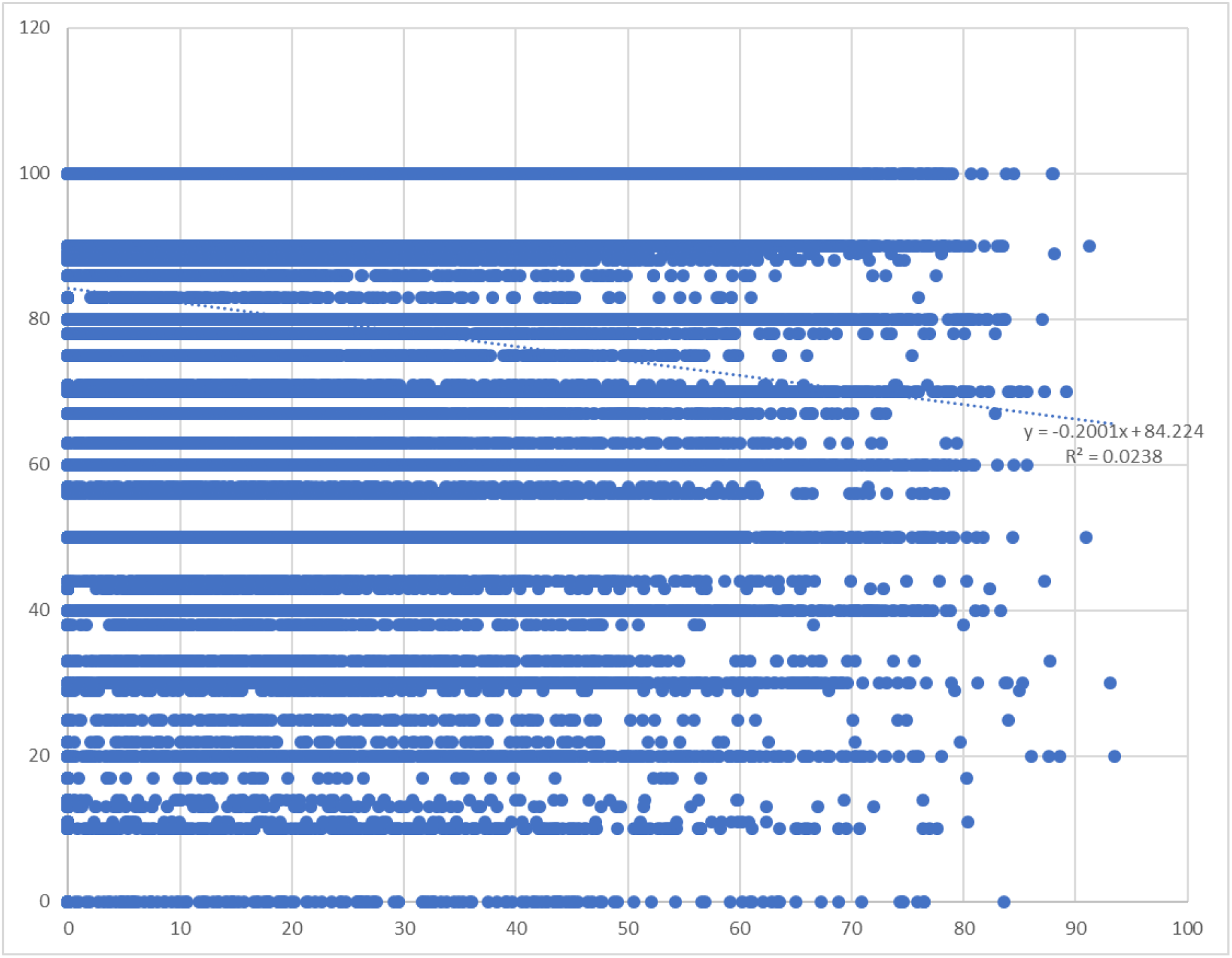
A scatter plot of all the coisolation interference and SPS%Accuracy measurement for all PSMS of the Fusion 1 files from PXD005361.

